# Sleep-insomnia superposition: opposing brain signatures of sleep in task-based and resting-state conditions

**DOI:** 10.1101/2023.05.13.540646

**Authors:** Mohamed Abdelhack, Peter Zhukovsky, Milos Milic, Shreyas Harita, Michael Wainberg, Shreejoy J Tripathy, John D Griffiths, Sean L Hill, Daniel Felsky

## Abstract

Sleep and depression have a complex, bidirectional relationship, with sleep-associated alterations in brain dynamics and structure impacting a range of symptoms and cognitive abilities. Previous work describing these relationships has provided an incomplete picture by investigating only one or two types of sleep measures, depression, or neuroimaging modalities in parallel. We analyzed the correlations between task and resting-state brain-wide signatures of sleep, cognition, and depression in over 30,000 individuals. Neural signatures of insomnia and depression were negatively correlated with neural signatures of sleep duration in the task condition but positively correlated in the resting-state condition, showing that resting-state neural signatures of insomnia and depression resemble that of rested wakefulness. This was further supported by our finding of hypoconnectivity in task but hyperconnectivity in resting-state data in association with insomnia and depression This information disputes conventional assumptions about the neurofunctional manifestations of hyper– and hypo-somnia, and may explain inconsistent findings in the literature.

## Introduction

The relationships between sleep, neurocognitive processes, and depression are complex and fraught with paradoxes. Depressive symptoms are linked to both increases and decreases in sleep duration. At the same time, acute sleep deprivation has been shown to act as an effective antidepressant^1^. Neural correlates of sleep are also linked to both hyper– and hypo-activation in different task environments^2, 3^, and both undersleeping and oversleeping are associated with poorer cognition^4^. The majority of our current understanding of sleep has come from acute sleep deprivation experiments and clinical sleep disorders which cover only fringe conditions in comparison with the general population^2, 3, 5, 6^. This has produced a gap in our understanding of the role of sleep in mental and cognitive health and how sleep affects the brain under different conditions and cognitive loads.

Observational studies have shown that sleep deprivation is associated with the degradation of attention, working memory, reward and dopamine processing, emotion discrimination and expression, and hippocampal memory processing^3^. Neurobiologically, it has been associated with aberrant activity observed in the visual cortex^7–9^, frontoparietal regions^10^, and ventral and dorsal attention networks^11^. This indicates a possible role of sleep in visual cortical processing via top-down attentional circuits during cognitive task performance.

Sleep quantity and quality are also linked to symptoms of mental illness^12–14^, and the biological mechanisms of these links have been explored with neuroimaging. For example, insomnia and depression are bidirectionally related^5^, with lower-quality sleep being associated with negative thoughts through decreased connections in the amygdala^15^. In one study, insomnia, daytime dozing, and low sleep quality were associated with aberrant functional connectivity in many brain regions, especially the default mode network^2^; however, most neuroimaging studies have been underpowered and yielded heterogeneous results, leading to inconclusive evidence. One challenge to parsing these relationships is that, while depression is typically associated with symptoms of insomnia^5^, atypical depression is associated with hypersomnia^16^. Insomnia is hypothesized to be a result of hyperarousal states that causes cognitive fatigue and anxiety that could progress to depression^17, 18^, while the relationship between hypersomnia and depression is still an open question.

One prevalent limitation of most published neuroimaging studies of sleep is that they rely on data collected during acute sleep deprivation, which does not necessarily have the same effect as chronic sleep loss or low sleep quality^3^. Further, studies that have investigated primary insomnia and depression have used a wide variety of methodologies, resulting in heterogeneous findings^2^. Only a few studies also analyze neural representations with subjective and objective sleep measures of sleep^19–21^. In addition, in primary insomnia, objective sleep measures using polysomnography do not align with the subjective report of participants^22, 23^.

Some initial attempts toward decoding these complex relationships at the population level have been made. Cheng et al. (2018) investigated the neural associations of sleep quality and depression using the Human Connectome Project (HCP)^24^ and a subset of UK Biobank^25^, finding that the link between poor sleep and depressive symptoms was in part mediated by patterns of functional connectivity. One limitation of this study was that they used the overall Pittsburgh Sleep Quality Index (PSQI) as their measure of sleep^26^ and only assessed resting state data, thus missing the heterogeneity of sleep-related phenotypes manifest under different conditions. More recently, Fan et al. (2022) performed a systematic analysis of multiple sleep phenotypes in a subset of white-British population of the UK Biobank^27^. They analyzed resting-state and task-based fMRI data, diffusion tensor imaging, and cardiac MRI data, testing their independent relationships with self-reported sleep data and other environment and mental health variables. They identified resting-state brain activation as a predictor of self-reported insomnia and narcolepsy. They also found no significant associations between task-based fMRI features and most sleep phenotypes. Crucially, this study was limited to self-reported sleep measures, and it did not investigate the relationships between neural signatures of tested phenotypes. Another study also investigated the associations of sleep phenotypes in association with obesity, cardiometabolic conditions, brain structure, and cognition but did not account for brain activity^28^.

To address these gaps, we perform a multi-step analysis of two independent cohorts and examine previously published analyses through a new lens. First, we map the associations of sleep quality – measured subjectively by self-report and objectively by accelerometry – with both task-based and resting-state measures of brain function in the UK Biobank^29, 30^. We then test the correlations between brain-wide patterns of associations with cognitive function and depressive symptoms, finding seemingly contradictory patterns of resting-state and task-based activation in response to poor sleep. Our analyses provide insights into shared mechanisms of the heterogeneity in depression symptoms and how they connect neurobiologically with sleep patterns. This could eventually lead to a better comprehension of the symptomology of depression in line with sleep patterns with the potential for more targeted therapies.

## Results

### Self-reported and accelerometer-based measures of sleep are weakly correlated

**Figure 1** summarizes the analyses performed in our study. We first quantified the pairwise phenotypic partial correlations between our five behavioral measures: sleep quality measured by an accelerometer (duration of longest sleep bout), self-reported sleeplessness/insomnia frequency, self-reported daytime dozing frequency, cognitive ability measured by a symbol-digit substitution task^31^, and subclinical depression symptoms measured by the PHQ-2^32^ with age, sex, study site, ethnicity, socioeconomic status, the difference between the time of accelerometer measurement and assessment center visit, and education as covariates. Accelerometer-measured sleep quality was weakly correlated with cognitive performance (r=0.036; *p*=5.39×10^-3^) while depressive symptoms were correlated with self-reported insomnia (r=0.15; *p*=5.64×10^-63^), both in positive directions (**Figure 1A**). Self-reported insomnia and daytime dozing frequencies were also positively correlated, though the magnitude of this correlation was similarly very small, with only 0.7% of variance explained (r=0.081; *p*=2.25×10^-20^). As expected, the accelerometer-measured duration of longest sleep bout had negative correlations with both self-reported insomnia (r=-0.072; *p*=2.21×10^-15^) and self-reported daytime dozing (r=-0.11; *p*=1.29×10^-35^), again with very small effect sizes.

**Figure 1:**
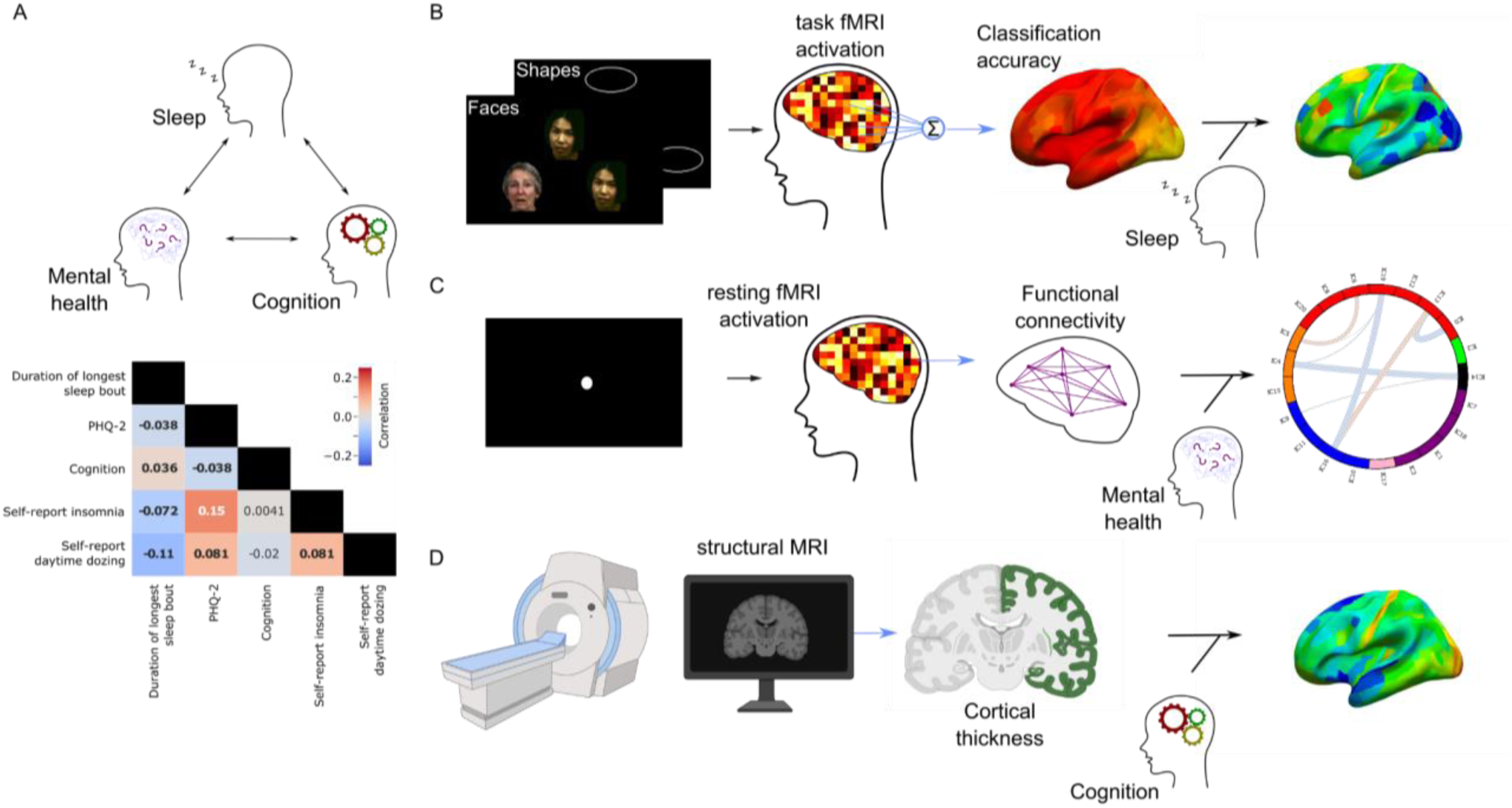
Study Summary. **A** shows the partial correlation map between the tested phenotypes of sleep (duration of longest sleep bout, self-reported insomnia, and self-reported daytime dozing), depression symptoms (PHQ-2 score), and cognition (bolded numbers are correlations significantly different from zero). **B** shows the task fMRI experiment, multivariate pattern analysis conducted, and the subsequent linear modeling of the classification accuracy with the phenotypes to build a cortical map of associations (stimulus images obtained by permission from Prof. Deanna Barch). **C** shows the resting-state data collection protocol, the calculation of functional connectivity, and the linear modeling to produce a connectivity association map. **D** shows the process for obtaining cortical thickness and linear modeling with phenotypes to generate a brain map similar to B.

### Multimodal neural associations with sleep, depression, and cognition

Having established phenotypic correlations between our measures, we first built a brain map of each phenotype using task-based fMRI (**Figure 1B**). We fit multivariate classification models^33, 34^ using support vector machines (SVM) to classify face and shape trials regardless of task performance. Models from all regions were able to significantly perform above the 50% chance level, however, classifiers using voxels from visual areas were the most accurate (**Figure S2**). We carried forward classification accuracies from each region as a proxy for its cortical activation in response to the visual stimuli. We then measured the association of this activation proxy with our phenotypes of interest using ordinary least squares (OLS) regression.

Our measure of task-based brain activation showed significant associations with accelerometer-measured sleep duration, depressive symptoms, and cognitive scores in predominantly visual regions as well as higher multimodal regions in the parietal cortex (**Figure 2A**). Cognition also showed significant associations across frontal regions while depressive symptom associations were more global and diffuse (**Figure 2B**). Longer sleep bouts were associated with a higher decoding accuracy (stronger multivariate cortical signal), primarily in lateral occipital regions (**Figure 2B, S4**). These are intermediate processing areas that feed into the ventral stream of vision. Higher regions along the ventral stream showed no significant associations with accelerometer-measured sleep. Depressive symptoms showed significant associations across regions spanning the whole cortex, where higher symptom scores were associated with lower decoding accuracies (**Figure 2B**). The strongest associations were observed in the visual areas, particularly the high-level face-selective and intermediate visual areas (**Figure S4**). Higher cognitive scores corresponded to higher decoding accuracy which overlapped with depressive symptoms score effects in visual cortex and prefrontal cortex (**Figure 2B,C**). The latter three phenotypes all had overlapping significant associations in multimodal superior parietal regions (**Figure 2C**). These areas are responsible for higher level visual processing of orientation and location as well as motor planning which is reasonable given the nature of the task involving visual recognition and motor action (button pressing). Self-reported insomnia frequency showed no significant effect on the neural coding of visual tasks except in one region in the prefrontal cortex. Self-reported daytime dozing frequency showed no significant associations in any region.

**Figure 2:**
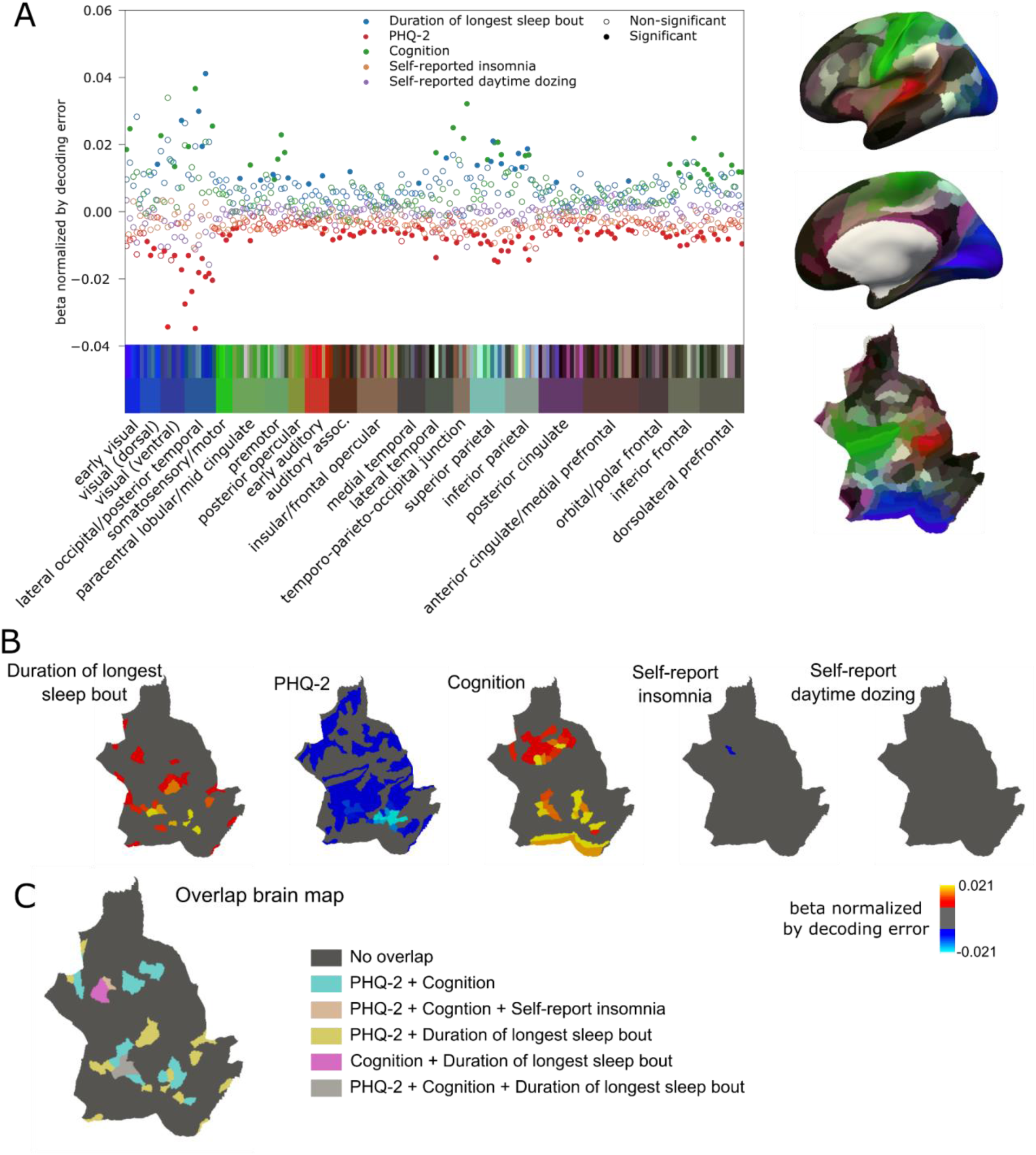
Results of the associations from the task-based fMRI data. **A)** summarizes the overall beta values of the models normalized by decoding accuracy. The regions are organized and color-coded according to their groupings in the human connectome project^35^ where the region color maps are shown on the right. **B)** shows the brain region maps with significant associations color-coded by the beta values normalized by decoding accuracy for each of the phenotype models. **C)** shows the overlap between regions that showed significant associations with more than one phenotype.

Following task-based analyses, we investigated the associations of resting-state data with our target phenotypes. We first analyzed associations of functional connectivity of independent components across the brain (**Figure 1C**), observing many significant associations with accelerometer-measured duration of longest sleep bout that spanned many circuits (**Figure 3A**). Daytime dozing showed a similar association pattern with opposite directions of effect due to the inverted scale of the two measures. Insomnia, depressive symptoms, and cognition were associated with only a few circuits and showed little overlap (**Figure S5**). From these associations, we selected one to probe in more detail using seed-based connectivity analysis.

**Figure 3:**
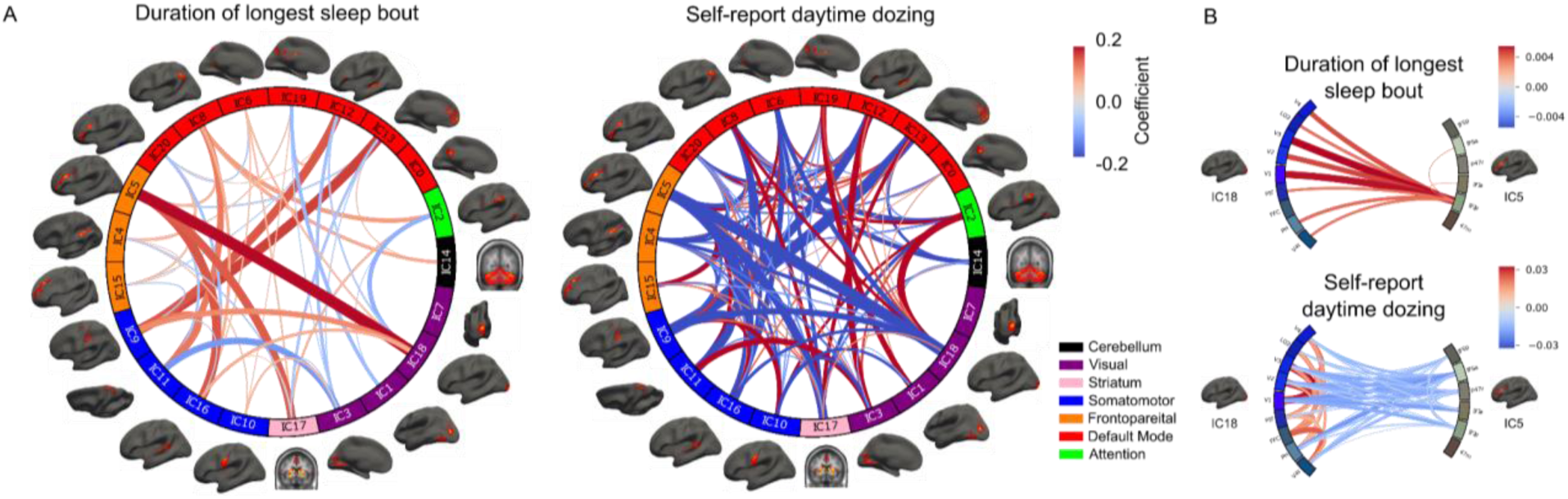
Resting-state connectivity associations results with the sleep phenotypes. **A)** shows the functional connectivity associations for the accelerometer-measured duration of longest sleep bout and self-reported daytime dozing. The different components are grouped and color-coded based on the Yeo 7 Networks^36^. **B)** zooms in on the associations between IC5 and IC18 showing seed-based correlation associations between different regions belonging to the components of interest.

Specifically, we investigated the connection between IC5 and IC18 as this could point to the reasons for the degraded signal in the task fMRI condition. The functional connectivity pattern between these two independent components was positively correlated with duration of longest sleep bout and negatively correlated with daytime dozing. We investigated the regions belonging to these two components by selecting the regions that mark higher than the 98th percentile of the component activation. Seed-based results showed a positive association with duration of longest sleep bout and negative association with daytime dozing at the connection level between the posterior side of the inferior frontal junction (IFJp) and almost all the occipital regions in IC18. This points to a positive association of sleep bout length with the connectivity between the frontal attentional areas and the intermediate visual regions. Complete associations of seed-based correlations are also shown in **Figure 6**.

Finally, we investigated the association of each phenotype with cortical thickness (**Figure 1D**). Measured duration of longest sleep bout, depressive symptoms, and cognition all showed significant and diffuse associations but with strongest overlap along the auditory, insular, and temporal regions (**Figure S7A**). Frequency of daytime dozing showed a sparse pattern that spanned many of the same regions. The results showed that higher cortical thickness was associated with longest continuous sleep (accelerometer-measured), less frequent depressive symptoms, higher cognitive score, and lower frequency of daytime dozing in almost all brain regions This pattern did not hold for the primary visual cortex (V1) and early visual cortex (V2 and V4). Self-reported insomnia did not show any significant association with cortical thickness values.

### Correlations of neural signatures of sleep, depression, and cognition show opposite relationships under task-activated vs. resting conditions

To quantify the similarity in brain-wide patterns of task-based association between phenotypes, we performed pairwise Pearson correlations between each set of association statistics in the task and resting state conditions. For the task-based condition, in directional agreement with our observed phenotypic correlations (**Figure 1A**), the neural signature of accelerometer-measured duration of longest sleep bout was negatively correlated with those for depressive symptoms (r=-0.63; *p*=5.07×10^-21^), frequency of insomnia (r=-0.14; *p*=0.04), and frequency of daytime dozing (r=-0.64; *p*=1.00×10^-21^). The neural signature for depressive symptoms showed positive correlations with those for both frequency of insomnia (r=0.17; *p*=0.03) and daytime dozing (r=0.64; *p*=2.36×10^-22^), indicating similar effects across the cortex despite the latter two phenotypes showing almost no significant independent associations (**Figure 4A**).

**Figure 4:**
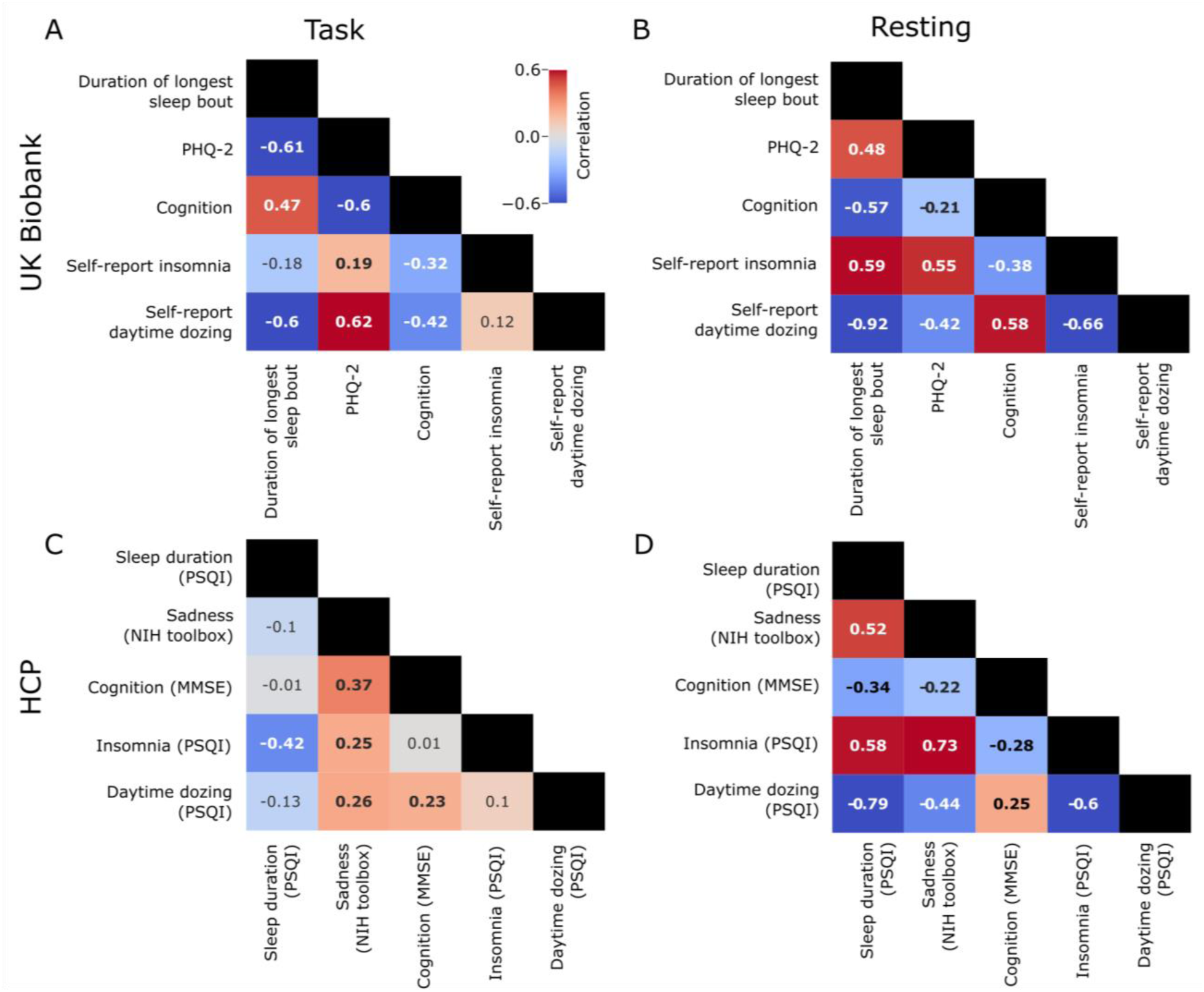
Task and resting conditions show a discrepancy in neural association correlations of sleep, cognition, and depression across two datasets. **A)** shows the pairwise correlation values between coefficients from each phenotype model of task-based activations across all brain regions in the UK Biobank. **B)** shows the pairwise correlation values between coefficients from each phenotype model of resting-state activations across all brain regions of the UK Biobank. **C)** task-based pairwise correlation measurement similar to **A** but for the HCP dataset. **D)** resting state pairwise correlation measurement similar to **B** but for the HCP dataset. Bolded values are statistically significant.

Shifting to neural signatures in the resting state condition, a notable difference emerged. In contrast to the results from the task condition and from phenotypic correlations, there were nontrivial positive correlations between the neural signature for duration of longest sleep bout and those for both self-reported insomnia (r=0.59; *p*=2.71×10^-21^) and depressive symptoms (r=0.48; *p*=1.87×10^-13^; **Figure 4B**). This indicated a similarity between the functional connectivity changes associated with longer continuous sleep, higher frequency of insomnia, and more depressive symptoms – which is counterintuitive. This pattern was similar, with the daytime dozing functional connectivity also showing negative correlations. To confirm the validity of these results, we retrieved independently modeled associations from Fan et al.^27^ between self-reported insomnia, daytime dozing, and sleep duration (data retrieved at http://www.ig4sleep.org/) and performed the same correlational analyses. Reassuringly, we found nearly identical patterns of correlations between effects (**Table S4**); self-reported insomnia and insomnia had a correlation coefficient of –0.66 (compared to –0.59 in our analysis). While they did not test accelerometer-measured duration of longest sleep bout, the results from self-reported sleep duration were consistent (correlation with self-reported insomnia=0.58, with daytime dozing=-0.86). To further confirm these findings, we performed a similar analysis on the independent HCP dataset, which included self-reported sleep measurements using PSQI^26^, sadness (proxy for depression) measured using the NIH toolbox^37^, and cognition measured by the Mini-Mental State Examination (MMSE)^38^. Results for both task-based and resting-state data were largely in agreement, with the exception of neural signatures for cognition measures (**Figure 4C, D**). Correlations between the associations of anatomical models were largely consistent with those from the task fMRI experiment (**Figure S7C**).

### Discrepant task-activated and resting fMRI signatures of sleep are partly reconciled by varying sleep duration

In order to investigate the counterintuitive yet durable positive correlation of insomnia and depression with longer sleep in resting state, we developed two hypotheses to explain it: 1) a subset of individuals reporting higher levels of depressive symptoms drive the discrepancy due to the fact that both oversleeping and insomnia are possible symptoms of depression, and 2) individuals with insomnia and depression symptoms possess resting-state neural patterns that resemble those with long sleep resulting in a hyperattentive state, preventing them from sleeping.

To test the first hypothesis, we split the participants by their depression symptoms into those who have a score of 3 or more as the depressed group and those who have a score less than 3 as the not-depressed group ^32^. We fitted the models for each group again and the same correlation patterns between phenotypes persisted in the not-depressed group. In the depressed group, the insomnia and duration of longest sleep bout correlation disappeared (**Figure 5A**). This could be a factor resulting from the fact that the depressed group was small, with only 944 participants vs. the remaining 29,918.

**Figure 5:**
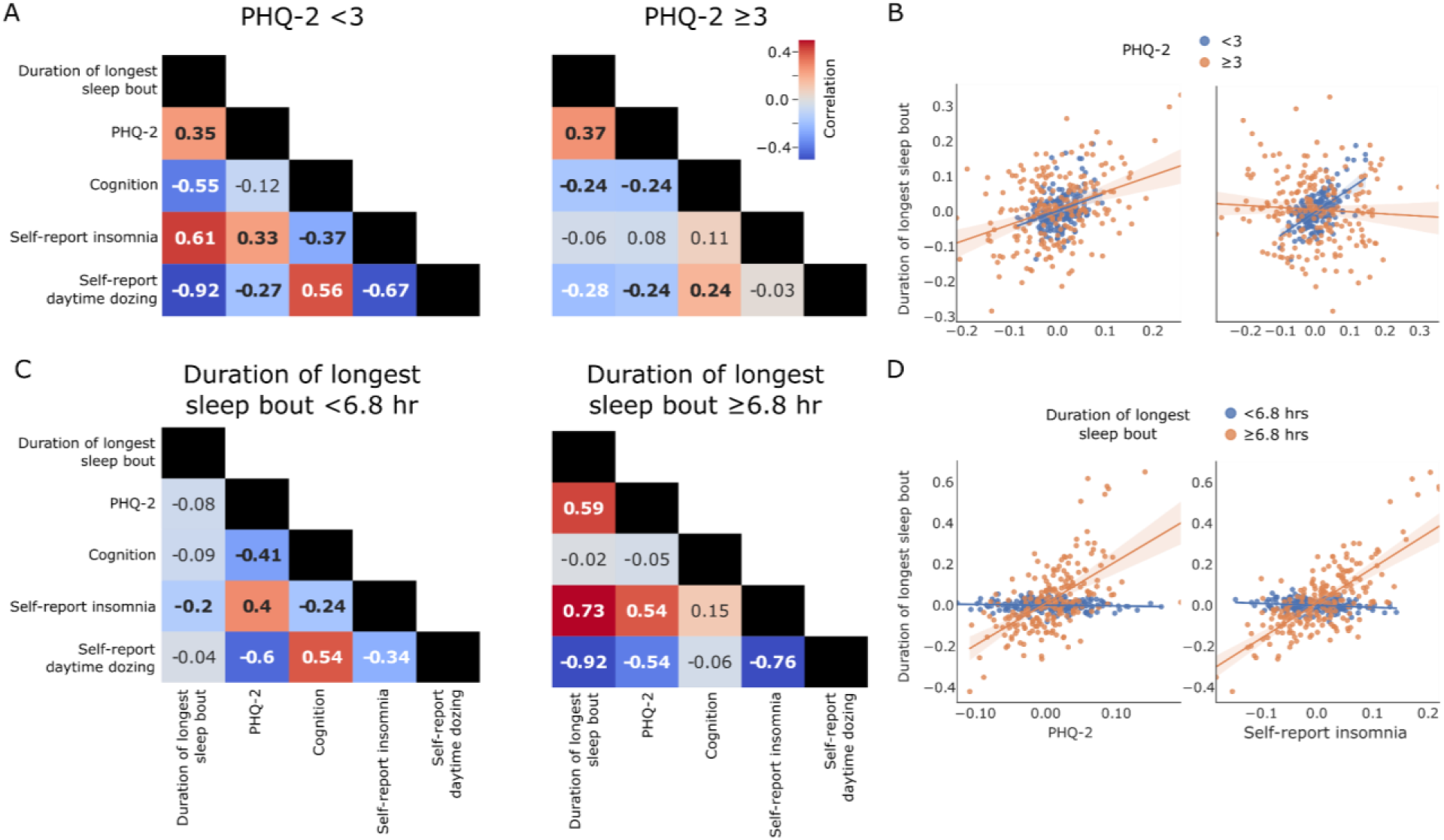
Correlation values of the coefficients of the linear models of functional connectivity values split by PHQ-2 scores and duration of longest sleep bout. **A** shows the pairwise correlation values for the coefficients of the models split by the PHQ-2 score representing depressed and non-depressed groups. **B** shows a scatter plot with the line of fit between the coefficients of the models for two pairs of phenotypes (duration of longest sleep bout and PHQ-2; duration of longest sleep bout and self-repot insomnia). **C** shows the pairwise correlation values for the coefficients of the models split by the duration of longest sleep bout. Bolded values are statistically significant. **D** shows a scatter plot with the line of fit between the coefficients of the models for two pairs of phenotypes (duration of longest sleep bout and PHQ-2; duration of longest sleep bout and self-repot insomnia).

We then split the participants into approximately equal groups split by the duration of longest sleep bout median value (greater or less than 6.8 hours). Individuals with an average of less than 6.8 measured hours of continuous sleep were labeled “short sleepers”, and those with an average of greater than or equal to 6.8 hours were labeled as “long sleepers” (**Figure 5C**). The positive correlation between sleep duration and both self-reported insomnia and depressive symptoms persisted only within the long sleepers (insomnia: r=-0.73; *p*=4.41×10^-36^; PHQ-2: r=-0.59; *p*=3.34×10^-21^). In the short sleeper group, we observed no significant correlation between neural signatures of sleep duration and that for PHQ-2 (r=-0.079; *p*=0.26), and we found a significant negative correlation of sleep duration with self-reported insomnia (r=-0.20; *p*=0.003). The positive correlation between signatures of depressive symptoms and self-reported insomnia persisted in both short and long sleepers. This result implies that sleep, when measured in “long sleepers”, relates to functional connectivity values that change in a pattern similar to increasing symptoms of depression and frequency of insomnia. However, significant negative correlations with the daytime dozing measure persisted in both groups but the negative correlation between duration of longest sleep bout and daytime dozing did not reach statistical significance.

### Brain regions are hyperconnected under the resting condition with depression and insomnia but hypoconnected during the task condition

In the previous section, we showed similar resting state patterns between depression and insomnia and duration of longest sleep bout. This was in contrast to the results from the task-based data. In order to investigate the directionality of associations of the neural connectivity patterns giving rise to this discrepancy, we compared global connectivity patterns across resting and task conditions. We calculated the representation connectivity patterns for the task condition and used seed-based connectivity from the previous analysis. We then modeled associations with our five sleep, depression, and cognition phenotypes across all pairs of brain regions as well as an aggregate brain-wide average connectivity measure (**Figure 6A,C**). We also modeled the association of the network-specific average connectivity (**Figure 6B**) in order to investigate intra– and inter-network connectivity changes. Results show that, for the task condition, there is a predominantly negative association between representational connectivity and depressive symptoms and self-reported insomnia (**Figure 6A**) suggesting hypoconnectivity association with these phenotypes. However, in the resting condition, the associations were mostly positive, suggesting hyperconnectivity. Self-reported daytime dozing showed a strongly negative association suggesting a strong hypoactivation in the resting condition. These results are also consistent with the correlation results (**Figure 4**). Results from the network-wise connectivities show a significant positive association between accelerometer-measured duration of longest sleep bout and the default mode network (DMN) inter-connectivity (**Figure 6B**). This significant association was also observed in self-reported insomnia but was accompanied by another significant association between the DMN and frontoparietal network (FPN). Similarly, depressive symptoms showed a significant positive association with the DMN-FPN connectivity but not within the DMN.

**Figure 6:**
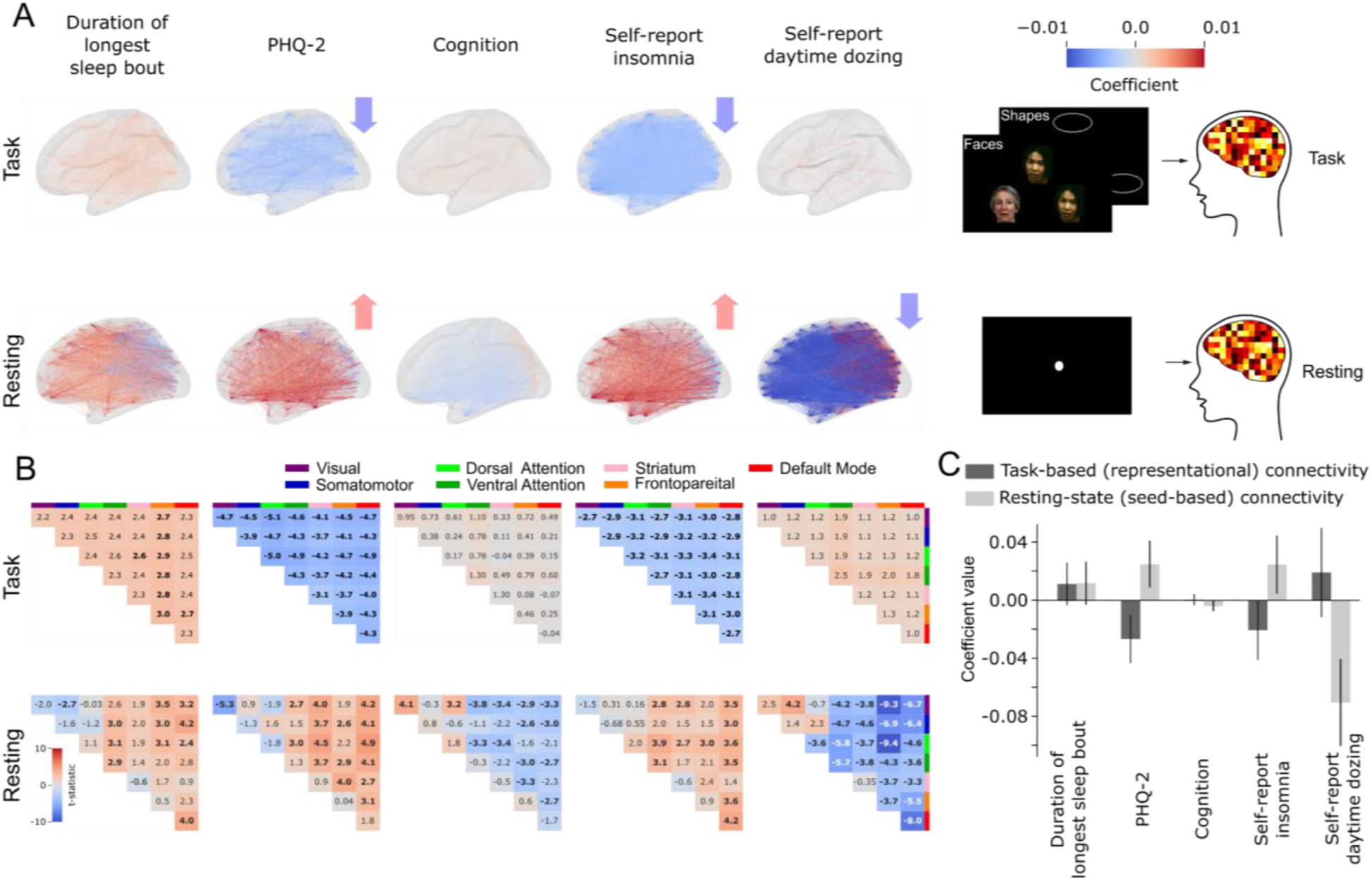
Brain regions are hyperconnected with PHQ-2 and insomnia in resting condition but hypoconnected in task condition. **A** shows the significant brain representational and functional connectivity associations with the five phenotypes for each connection between HCP180 regions. **B** Network-wise representational (upper) and functional (lower) connectivity associations (model t-statistic) with the different phenotypes. Bolded associations are statistically significant. **C** Global mean connectivity associations with the five phenotypes. Error bars represent 95% confidence intervals after correcting for multiple comparisons over 5 measures.

## Discussion

We observed a striking and consistent contrast between the neural representations of objectively-measured and self-reported sleep. Specifically, brain-wide resting state fMRI signatures of long accelerometer-measured sleep were the same as those of higher self-reported frequency of insomnia and depressive symptoms. This seemingly paradoxical result was replicated using summary statistics from a previously published study and in independent analyses of the HCP dataset. Under task conditions, these correlations were inverted. This discrepancy was partially reconciled by showing that the positive correlations in resting state data persisted only for individuals with sleep durations measured on average longer than 6.8 hours. Additionally, brain-wide mean connectivity increased with insomnia and depression at resting state but decreased under the task condition. Our findings may explain heterogeneity in existing literature on the neural signatures of sleep and depression, and shed new light on the specific circuits responsible for the connections between sleep, depression, and cognition.

Our task-based analyses relied on a measure of signal-based decoding of task trials using machine learning. Superior parietal regions showed significant associations with the duration of longest sleep bout, depressive symptoms, and cognition. Insomnia and dozing showed few significant associations, in line with previous univariate analyses on the same data^27^. Conversely, objective sleep measure revealed associations with neural data sensitivity to neural activity changes in comparison to self-reporting. Duration of longest sleep bout had additional associations with intermediate visual areas at the lateral occipital junction, with better sleep being associated with higher multivariate activation; these areas are responsible for shape detection^39^.

In addition, resting state results revealed widespread associations similar to Fan et al. 2022^27^, especially for daytime dozing. We found that functional connectivity between the frontoparietal network and in particular the posterior inferior frontal junction (IFJp) and lateral occipital regions was positively associated with duration of longest sleep bout. IFJp is known to be responsible for top-down attention^40, 41^. This suggests an effect of sleep on the top-down visual attention connections leading to degraded visual processing. It is known that top-down attention can modulate visual cortex activation patterns^42^ and thus any impairment in this connection could impair visual function. This effect was reported previously in patients with primary insomnia^43, 44^. Previous experiments of sleep deprivation have shown a decreased connectivity between frontal and parietal regions with the visual cortex^45–47^ and a decrease in activation of the visual cortex^7–9^ that was reversible using trans magnetic stimulation^48, 49^. It challenges the results from previous sleep deprivation studies that report a decrease in attention signal at the source at the dorsolateral prefrontal cortex^45–47, 50–53^ suggesting instead a connectivity impairment. These studies relied mostly on acute sleep deprivation that could lead to transient impairment in cognition as opposed to sustained low sleep quality where connectivity becomes impaired as a result of sustained low attentional signal from the source.

Our central finding was that functional connectivity signatures were positively correlated between longer bouts of accelerometer-measured sleep and both frequency of self-reported insomnia and greater depressive symptoms. This correlation remained in both strata of high and low depressive symptoms, but only persisted in long sleepers when the population was stratified by longest duration of sleep bout. The positive correlation between long sleep and depressive symptoms could in part explain an atypical presentation of depression symptoms: hypersomnia^16^. The positive correlation of long sleep with insomnia could have two explanations: one is that the resting-state signal of a person with a higher frequency of insomnia resembles that of a rested wakefulness state thus preventing them from falling asleep and keeping them in a hyperarousal state^5, 13, 17, 18^. Results from sleep EEG suggest that during sleep, signals resemble a hyperarousal state decreasing the quality of sleep in insomnia^54^. Another possibility is that the objective measure of sleep by accelerometry is not capturing the objective sensation of sleep quality which is reported by primary insomnia patients and polysomnography measurements^22, 23^. However, we believe that the first explanation is more likely given the phenomenon of contradictory subjective and objective sleep measure results was observed in polysomnography but not accelerometry measures^19–21, 55–57^ and that our results were reproduced in the HCP dataset where sleep duration was self-reported^24^. This pattern was also reproducible through analyzing publicly available coefficients from an independent analysis of UK Biobank^27^.

In our population, the groups of short sleepers (duration of longest sleep bout < 6.8 hours) showed an inverted association with insomnia which is reasonable but it signals that insomnia neural signature is multimodal resembling both short and long sleep. There was no significant association between duration of longest sleep bout and PHQ-2 in that group. The positive associations between duration of longest sleep bout remained consistent between depressed and not-depressed groups while insomnia association was insignificant for the depressed group.

The non-depressed group showed identical associations with the whole cohort which could be explained that the non-depressed group represented the majority of the cohort. Brain-wide mean connectivity results revealed that insomnia and depression are associated with hypoconnectivity in the task condition and hyperconnectivity during the resting condition. Previous studies have found similar results of hypoactivation in primary insomnia for task-based fMRI^58, 59^ while resting state connectivity results in the literature were mixed^44, 60^. For depression, hyperconnectivity was observed in various networks for resting conditions^61, 62^. In addition, sleep state is associated with a breakdown of cortical effective connectivity^63, 64^ so insomnia being associated with hyperconnectivity in the resting state could signal a reverse effect.

Our study has several limitations. First, we studied a general population sample with only a small minority of participants diagnosed with depression, insomnia, dementia, or narcolepsy. Therefore, our findings may not extend to clinical populations with severe impairments and symptoms. Second, while the results of our analyses in UK Biobank and HCP were largely consistent, task signatures of cognition with those for other phenotypes were not entirely consistent. This may have been due to differences in cognitive measures in these two cohorts. The measure of cognition used in the UK Biobank analysis was a word-symbol matching task where changes in performance could indicate cognitive decline. In HCP analyses, we utilized the available test for cognitive decline, the MMSE, but these two measures might not capture the heterogeneity of brain functions that show dysfunction with cognitive decline. This is especially evident in our UK Biobank analysis where the cognitive measure was derived from a single task. Similarly, the results we obtained from our task-based fMRI study might not necessarily generalize to tasks other than face-shape matching which could limit the conclusions of this analysis.

Our results show that longer uninterrupted sleep is related to the strength of sensory and cognitive processing in vision areas, possibly due to the increased top-down attention recruitment. Additionally, we counterintuitively found similarities in resting state activity among people with insomnia, long sleep, and depression symptoms which could signal hyperarousal in resting state activity. This hyperarousal could increase the possibility of cognitive fatigue that may end up causing a reduction in task-based activation. This persistent fatigue could give rise to depressive symptoms with daytime dozing acting as a compensatory mechanism, and may partly explain the success of sleep deprivation as a therapy for depression^1^. It also highlights the heterogeneity of sleep quality factors where previous studies showed hyperconnectivity to be associated with poor sleep with dorsolateral prefrontal cortex, cuneus, and orbitofrontal cortex mediating the relationship with depression ^25^. That previous work, however, used the overall PSQI sleep score as a marker for poor sleep. We showed here that within the same HCP cohort that they used, different components of the PSQI score have different neural signatures.

Our study highlights the importance of investigating the multimodal signature of phenotypes to understand their diverse manifestations that could give rise to similar symptoms. Our results are supported by a large sample size of over 30,000 participants from the UK Biobank and over 800 from HCP study. The sheer size of these datasets also allows for studying more brain-wide associations with reproducible quality and relatively accurate effect sizes^65^. We uncover a phenomenon of brain-wide similarities between sleep quality, insomnia, and depressive symptoms that could guide advancing clinical practice to investigate more fine-grained details of sleep habits to guide the optimal care plans all while concurrently tracking the cognitive load of patients.

## Methods

### Software

We utilized FreeSurfer and FSL tools for brain region parcellation and label transformation as well as for cortical thickness measurements and seed-based correlation analysis and higher-level modeling of its results. We used python 3.6 for subsequent analyses with Brain Decoder Toolbox 2 for brain region data extraction, scikit-learn for SVM classifier construction, statsmodels for OLS model creation. For plotting and visualization, we used the libraries seaborn^66^ and mne-connectivity.

### Dataset

Data was obtained from UK Biobank^29, 30^ application #61530. We collected data for the functional magnetic resonance imaging for the resting state and task-based paradigms as well as the anatomical data. We also utilized the task data from E-Prime software (software data) to characterize the task-based runs. For sleep data, we obtained both data from the self-report sleep quality measures collected at the same imaging instance and from wrist-based accelerometers, which were worn over a 7-day period and used for extracting quantitative measurements of sleep quality^14^. Other psychiatric (PHQ-2) and cognitive measures (symbol digit substitution task) were collected from the self-reported mental health questionnaires and cognitive test results^31^ conducted at the same imaging instance. **Table S1** indicates the variable codes from UK Biobank and the number of valid subjects extracted for each data modality.

Covariates were extracted from the demographics data in UK Biobank (sex, age, socioeconomic status, ethnicity, and education level) and the measurement-specific factors (difference in time between accelerometry measurement and brain image acquisition, head motion, face-shape task performance, and measurement site). To maximize the number of participants and strengthen statistical power in each association analysis, we included all participants with an available measurement for each phenotype independently rather than investigating only the participants with all valid measures (**Figure S1**). This led to different numbers of participants for each phenotype measurement (**Table S2**).

### Phenotypic correlation analysis

We measured the pairwise phenotypic partial correlations between five output parameters: sleep quality measured by an accelerometer (duration of longest sleep bout), self-reported sleeplessness/insomnia frequency, self-reported daytime dozing frequency, cognitive ability measured by number of correct matches in a symbol-digit substitution task^31^, and subclinical depression score measured by the PHQ-2 scale^32^ with age, sex, study site, ethnicity, socioeconomic status, difference between time of accelerometer measurement and assessment center visit (only for the accelerometer output parameter), and education level as covariates. We calculated confidence intervals and significance by the 99% confidence intervals to correct for multiple comparisons (0.05 significance level over five outputs).

### ROI-based analysis

Regions of interest for the multivariate pattern analyses were constructed using the predefined cortical parcellations from the Human Connectome Project^35^. We combined the bilateral regions of interest resulting in 180 parcellations. The labels from the HCP parcellation were transformed using FreeSurfer software^67^ from the fsaverage subject cortical surface to each subject’s surface in the dataset. Labels were then transformed into the volume space of the fMRI data for each of the resting state and task-based paradigms.

### Task fMRI analysis

The task fMRI experiment in UK Biobank data comprised a modified version of the face-shape matching task^68, 69^. In this task subjects viewed a central cue stimulus accompanied by two stimuli on the right and the left with one of them matching the central cue. Subjects were tasked to press a button identifying which of the two stimuli is the one matching the central cue. The trials contained either human faces or 2D shapes (circle, horizontal ellipse, and vertical ellipse). In order to perform brain-wide association analysis with the task-based fMRI data, we built multivariate classification models^33, 34^ using support vector machines (SVM) to classify the face and shape trials regardless of subject’s performance. Models were created for each region of interest where regions were delineated according to the human connectome project (HCP) parcellation^69^. We carried forward the classification accuracies from each region as a proxy for its cortical activation in response to the visual stimuli. We then measured the association of classification accuracy with our phenotypes of interest using ordinary least squares (OLS) regression models. We created ordinary least square models relating the classification accuracy of each region and sleep efficiency. We also added the relevant covariates to the model (sex, age, imaging site, head motion, socioeconomic status, education level, ethnicity, task performance accuracy mean, task response time mean, task response time standard deviation, sex and age interaction, and accelerometry time relative to brain acquisition). To correct for multiple comparisons, we adjusted the p-values for the false discovery rate using the Benjamini/Hochberg method. We then divided the resulting model coefficients by the classification error to up-weight regions with voxels most responsive to the stimuli.

### Multivariate pattern analysis

We utilized the readily preprocessed task-based fMRI data from UK Biobank to create classifiers between faces and shapes for each brain region. Time series from each region was extracted using Brain Decoder Toolbox 2 for Python (https://github.com/KamitaniLab/bdpy). We then applied further preprocessing to the data where the data volumes were shifted by 5 volumes (3.675 seconds) to compensate for the hemodynamic delay. Data was then filtered to remove the slow signal shift along the run, and then samples were normalized by the mean value to extract the percent signal change. We then averaged the samples belonging to the same classification category within each block to improve the signal-to-noise ratio. Finally, the data points without stimulus were removed and the samples were then randomized. We ended up with 60 data points for the classifier which were then randomized and divided into training and test datasets in a 6-fold cross-validation scheme. For each fold, we trained a binary support vector machine classifier with a linear kernel to classify the faces and shapes. The mean classification accuracy from each region was then calculated and utilized as a proxy for the strength of encoding of stimuli in this brain region.

### Representational connectivity analysis

We extracted and preprocessed the task fMRI data in a similar fashion as in the MVPA analysis. We then divided the stimuli into 7 different categories based on the content of stimuli with three categories representing shapes (circle, horizontal ellipse, and vertical ellipse) and four representing faces (male, female, angry, and fearful faces). The voxel data for each of these conditions were then averaged creating a vector of voxel data for each region. We then computed the representational dissimilarity matrix (RDM)^70^ for each region. To calculate representational connectivity, we conducted a second-order similarity analysis between region pairs by calculating the Pearson correlation coefficient between the lower triangles of their RDMs.

### Cortical thickness measurement

Cortical thickness was measured for each brain region using Freesurfer software anatomical statistics measurement tools using the FreeSurfer reconstructed brain anatomy images provided by UK Biobank. We then created ordinary least squares models relating cortical thickness data to sleep efficiency and relevant covariates (sex, age, socioeconomic status, education level, ethnicity, imaging site, sex and age interaction, accelerometry time relative to brain acquisition). To correct for multiple comparison, we adjusted the p-values for false discovery rate using the Benjamini/Hochberg method.

### Functional connectivity analysis

We extracted the readily-processed functional connectivity data based on full correlation from UK Biobank repository (variable code: 25750) and created ordinary least square models relating functional connectivity between each node (independent component) and sleep efficiency. We also added the relevant covariates to the model (sex, age, imaging site, head motion, socioeconomic status, education level, ethnicity, sex and age interaction, and accelerometry time relative to brain acquisition). To correct for multiple comparisons, we adjusted the *p*-values for multiple comparison using Bonferroni’s correction for five phenotypes and 21 independent components.

### Seed-based correlation analysis

In order to create more fine-grained connectivity patterns that also map to the same regions as the task-based fMRI, we ran a seed-based correlation analysis on each region using FSL dual regression tool^71^. We then divided the resulting correlation map into the HCP region space computing the mean over each region resulting in a 180 x 180 matrix of connectivity. We then normalized the rows of the matrix by the self-correlation values (diagonal of the matrix). Results were used to construct a higher-level model with sleep efficiency as the independent variable and the resting state covariates similar to the OLS models previously described in the functional connectivity analysis.

### Brain-wide mean connectivity analysis

We calculated brain-wide mean connectivity by averaging the seed-based connectivity across all node pairs from the HCP regions for the resting state data. For the task-based data we averaged the representational connectivity measures across all the regions. We then built OLS models for each mean connectivity value for each phenotype and calculated the model coefficients and confidence intervals based on a *p*-value of 0.01 based on Bonferroni correction for five phenotypes.

### Human connectome project data analysis

We extracted the HCP data from the young adult project^24, 72^. We extracted the data for the emotion task and the resting-state. For the emotion task, there were two runs for each subject with an identical task as that of the UK Biobank. We concatenated the data for these two runs and constructed the SVM models similar to the protocol used for UK Biobank. For the resting-state data, we utilized the already processed functional connectivity based on the full correlation between nodes defined by the group-ICA analysis.

For the phenotypes equivalent to those we analysed in the UK Biobank, we used the sleep parameters based on the PSQI sleep score^26^ as there was no objective sleep measures. We utilized the self-reported sleep duration as a proxy for the accelerometer-measured duration of longest sleep bout, the PSQI second component that relies on difficulty of falling asleep as a proxy for insomnia, and the answer to the question on trouble staying awake during daytime activities as a proxy for daytime dozing. For depression measure, we used the reported sadness score from the assessment of self-reported negative affect measure from the NIH toolbox^37^. For cognition, despite the HCP data containing cognitive test, the test score we utilized for the UK Biobank was not done for the HCP cohort. We utilized the Mini-Mental State Examination results as generic test for cognition^38^. The complete set of subjects with all imaging and behavioral phenotypes available was 807.

## Acknowledgments

The authors would like to thank the members of the Whole Person and Population Modelling group members for their valuable support, discussions, and feedback throughout the different stages of this study. We also thank Qing Chang for his help with data access and Prof. Colin Hawko and Dr. Erin Dickie for their helpful discussions over fMRI data processing methodology. We would also like to thank the participants in the UK Biobank and HCP studies for their contribution to open science. We also thank Prof. Deanna Barch for allowing us to use the stimulus images and the UK Biobank support and community for their help with the understanding of data.

## Author roles

Conceptualization: MA and DF

Method development: MA and DF

Data cleaning and analysis: MA, SH, and PZ

Visualization: MA and PZ

Supervision: DF, SLH, GG

Writing – Original Draft: MA and DF

Writing – review and editing: MA, DF, SLH, SH

## Data and Code Availability

UK Biobank data is available through an application process. More information about it is available through this URL: https://www.ukbiobank.ac.uk/enable-your-research/apply-for-access. HCP Young Adult data is freely available through this URL: https://www.humanconnectome.org/study/hcp-young-adult/data-releases. Codes to analyze this data are currently being organized for sharing and will be available on Github.

## References

1. Boland, E. M. et al. Meta-Analysis of the Antidepressant Effects of Acute Sleep Deprivation. J. Clin. Psychiatry 78, 893 (2017).

2. Fasiello, E. et al. Functional connectivity changes in insomnia disorder: A systematic review. Sleep Med. Rev. 61, 101569 (2022).

3. Krause, A. J. et al. The sleep-deprived human brain. Nat. Rev. Neurosci. 18, 404–418 (2017).

4. Tai, X. Y., Chen, C., Manohar, S. & Husain, M. Impact of sleep duration on executive function and brain structure. *Commun*. Biol. 5, 1–10 (2022).

5. Baglioni, C., Spiegelhalder, K., Lombardo, C. & Riemann, D. Sleep and emotions: A focus on insomnia. Sleep Med. Rev. 14, 227–238 (2010).

6. Fortier-Brochu, É., Beaulieu-Bonneau, S., Ivers, H. & Morin, C. M. Insomnia and daytime cognitive performance: A meta-analysis. Sleep Med. Rev. 16, 83–94 (2012).

7. Choo, W.-C., Lee, W.-W., Venkatraman, V., Sheu, F.-S. & Chee, M. W. L. Dissociation of cortical regions modulated by both working memory load and sleep deprivation and by sleep deprivation alone. NeuroImage 25, 579–587 (2005).

8. Chee, M. W. L. & Chuah, Y. M. L. Functional neuroimaging and behavioral correlates of capacity decline in visual short-term memory after sleep deprivation. Proc. Natl. Acad. Sci. 104, 9487–9492 (2007).

9. Habeck, C. et al. An event-related fMRI study of the neurobehavioral impact of sleep deprivation on performance of a delayed-match-to-sample task. *Cogn*. Brain Res. 18, 306– 321 (2004).

10. Li, G. et al. Magnetic resonance study on the brain structure and resting-state brain functional connectivity in primary insomnia patients. Medicine (Baltimore) 97, e11944 (2018).

11. Yeo, B. T. T., Tandi, J. & Chee, M. W. L. Functional connectivity during rested wakefulness predicts vulnerability to sleep deprivation. NeuroImage 111, 147–158 (2015).

12. Leistedt, S. J. J. et al. Altered sleep brain functional connectivity in acutely depressed patients. Hum. Brain Mapp. 30, 2207–2219 (2009).

13. Fernández-Mendoza, J. et al. Cognitive-Emotional Hyperarousal as a Premorbid Characteristic of Individuals Vulnerable to Insomnia. Psychosom. Med. 72, 397–403 (2010).

14. Wainberg, M. et al. Association of accelerometer-derived sleep measures with lifetime psychiatric diagnoses: A cross-sectional study of 89,205 participants from the UK Biobank. PLOS Med. 18, e1003782 (2021).

15. Motomura, Y. et al. Sleep Debt Elicits Negative Emotional Reaction through Diminished Amygdala-Anterior Cingulate Functional Connectivity. PLOS ONE 8, e56578 (2013).

16. Singh, T. & Williams, K. Atypical Depression. Psychiatry Edgmont 3, 33–39 (2006).

17. Riemann, D. et al. The hyperarousal model of insomnia: A review of the concept and its evidence. Sleep Med. Rev. 14, 19–31 (2010).

18. Bonnet, M. H. & Arand, D. L. Hyperarousal and insomnia: State of the science. Sleep Med. Rev. 14, 9–15 (2010).

19. Nie, X. et al. Functional connectivity of paired default mode network subregions in primary insomnia. Neuropsychiatr. Dis. Treat. 11, 3085–3093 (2015).

20. Liu, X., Zheng, J., Liu, B.-X. & Dai, X.-J. Altered connection properties of important network hubs may be neural risk factors for individuals with primary insomnia. Sci. Rep. 8, 5891 (2018).

21. Dai, X.-J. et al. Altered inter-hemispheric communication of default-mode and visual networks underlie etiology of primary insomnia. Brain Imaging Behav. 14, 1430–1444 (2020).

22. Feige, B. et al. Does REM sleep contribute to subjective wake time in primary insomnia? A comparison of polysomnographic and subjective sleep in 100 patients. J. Sleep Res. 17, 180–190 (2008).

23. Perlis, M. L., Giles, D. E., Mendelson, W. B., Bootzin, R. R. & Wyatt, J. K. Psychophysiological insomnia: the behavioural model and a neurocognitive perspective. J. Sleep Res. 6, 179–188 (1997).

24. Van Essen, D. C. et al. The Human Connectome Project: a data acquisition perspective. NeuroImage 62, 2222–2231 (2012).

25. Cheng, W., Rolls, E. T., Ruan, H. & Feng, J. Functional Connectivities in the Brain That Mediate the Association Between Depressive Problems and Sleep Quality. JAMA Psychiatry 75, 1052–1061 (2018).

26. Buysse, D. J., Reynolds, C. F., Monk, T. H., Berman, S. R. & Kupfer, D. J. The Pittsburgh sleep quality index: A new instrument for psychiatric practice and research. Psychiatry Res. 28, 193–213 (1989).

27. Fan, Z. et al. Mapping sleep’s phenotypic and genetic links to the brain and heart: a systematic analysis of multimodal brain and cardiac images in the UK Biobank. 2022.09.08.22279719 Preprint at https://doi.org/10.1101/2022.09.08.22279719 (2022).

28. Yu, J. et al. Associations between sleep-related symptoms, obesity, cardiometabolic conditions, brain structural alterations and cognition in the UK biobank. Sleep Med. 103, 41– 50 (2023).

29. Miller, K. L. et al. Multimodal population brain imaging in the UK Biobank prospective epidemiological study. Nat. Neurosci. 19, 1523–1536 (2016).

30. Alfaro-Almagro, F. et al. Image processing and Quality Control for the first 10,000 brain imaging datasets from UK Biobank. NeuroImage 166, 400–424 (2018).

31. Jaeger, J. Digit Symbol Substitution Test. J. Clin. Psychopharmacol. 38, 513–519 (2018).

32. Kroenke, K., Spitzer, R. L. & Williams, J. B. W. The Patient Health Questionnaire-2: Validity of a Two-Item Depression Screener. Med. Care 41, 1284–1292 (2003).

33. Haxby, J. V. et al. Distributed and Overlapping Representations of Faces and Objects in Ventral Temporal Cortex. Science 293, 2425–2430 (2001).

34. Kamitani, Y. & Tong, F. Decoding the visual and subjective contents of the human brain. Nat. Neurosci. 8, 679–685 (2005).

35. Glasser, M. F. et al. A multi-modal parcellation of human cerebral cortex. Nature 536, 171– 178 (2016).

36. Thomas Yeo, B. T., et al. The organization of the human cerebral cortex estimated by intrinsic functional connectivity. J. Neurophysiol. 106, 1125–1165 (2011).

37. Pilkonis, P. A. et al. Assessment of self-reported negative affect in the NIH Toolbox. Psychiatry Res. 206, 88–97 (2013).

38. Arevalo-Rodriguez, I. et al. Mini-Mental State Examination (MMSE) for the early detection of dementia in people with mild cognitive impairment (MCI). Cochrane Database Syst. Rev. 2021, CD010783 (2021).

39. Grill-Spector, K., Kourtzi, Z. & Kanwisher, N. The lateral occipital complex and its role in object recognition. Vision Res. 41, 1409–1422 (2001).

40. Bj, T.-R., Cl, A. & R, M. Functional dissociation of the inferior frontal junction from the dorsal attention network in top-down attentional control. J. Neurophysiol. 120, (2018).

41. Meyyappan, S., Rajan, A., Mangun, G. R. & Ding, M. Role of Inferior Frontal Junction (IFJ) in the Control of Feature versus Spatial Attention. J. Neurosci. 41, 8065–8074 (2021).

42. Horikawa, T. & Kamitani, Y. Attention modulates neural representation to render reconstructions according to subjective appearance. *Commun*. Biol. 5, 1–12 (2022).

43. Huang, S. et al. Regional impairment of intrinsic functional connectivity strength in patients with chronic primary insomnia. Neuropsychiatr. Dis. Treat. 13, 1449–1462 (2017).

44. Li, S. et al. Altered resting state connectivity in right side frontoparietal network in primary insomnia patients. Eur. Radiol. 28, 664–672 (2018).

45. Chee, M. W. L. & Tan, J. C. Lapsing when sleep deprived: Neural activation characteristics of resistant and vulnerable individuals. NeuroImage 51, 835–843 (2010).

46. Chee, M. W. L. et al. Effects of sleep deprivation on cortical activation during directed attention in the absence and presence of visual stimuli. NeuroImage 58, 595–604 (2011).

47. Chee, M. W. L., Tan, J. C., Parimal, S. & Zagorodnov, V. Sleep deprivation and its effects on object-selective attention. NeuroImage 49, 1903–1910 (2010).

48. Luber, B. et al. Remediation of Sleep-Deprivation–Induced Working Memory Impairment with fMRI-Guided Transcranial Magnetic Stimulation. Cereb. Cortex 18, 2077–2085 (2008).

49. Luber, B. et al. Extended Remediation of Sleep Deprived-Induced Working Memory Deficits Using fMRI-guided Transcranial Magnetic Stimulation. Sleep 36, 857–871 (2013).

50. Thomas, M. et al. Neural basis of alertness and cognitive performance impairments during sleepiness. I. Effects of 24 h of sleep deprivation on waking human regional brain activity. J. Sleep Res. 9, 335–352 (2000).

51. Chee, M. W. L. et al. Lapsing during Sleep Deprivation Is Associated with Distributed Changes in Brain Activation. J. Neurosci. 28, 5519–5528 (2008).

52. Czisch, M. et al. On the need of objective vigilance monitoring: Effects of sleep loss on target detection and task-negative activity using combined EEG/fMRI. Front. Neurol. 3, (2012).

53. Tomasi, D. et al. Impairment of Attentional Networks after 1 Night of Sleep Deprivation. Cereb. Cortex 19, 233–240 (2009).

54. Perlis, M. L., Merica, H., Smith, M. T. & Giles, D. E. Beta EEG activity and insomnia. Sleep Med. Rev. 5, 365–376 (2001).

55. Pace-Schott, E. F. et al. Resting state functional connectivity in primary insomnia, generalized anxiety disorder and controls. Psychiatry Res. Neuroimaging 265, 26–34 (2017).

56. Natale, V., Plazzi, G. & Martoni, M. Actigraphy in the Assessment of Insomnia: A Quantitative Approach. Sleep 32, 767–771 (2009).

57. Hauri, P. J. & Wisbey, J. Wrist Actigraphy in Insomnia. Sleep 15, 293–301 (1992).

58. Li, Y. et al. Abnormal Neural Network of Primary Insomnia: Evidence from Spatial Working Memory Task fMRI. Eur. Neurol. 75, 48–57 (2016).

59. Kay, D. B. & Buysse, D. J. Hyperarousal and Beyond: New Insights to the Pathophysiology of Insomnia Disorder through Functional Neuroimaging Studies. Brain Sci. 7, 23 (2017).

60. Schilbach, L. et al. Meta-Analytically Informed Network Analysis of Resting State fMRI Reveals Hyperconnectivity in an Introspective Socio-Affective Network in Depression. PLOS ONE 9, e94973 (2014).

61. Kaiser, R. H., Andrews-Hanna, J. R., Wager, T. D. & Pizzagalli, D. A. Large-Scale Network Dysfunction in Major Depressive Disorder: A Meta-analysis of Resting-State Functional Connectivity. JAMA Psychiatry 72, 603–611 (2015).

62. Zhu, Z. et al. Hyperconnectivity between the posterior cingulate and middle frontal and temporal gyrus in depression: Based on functional connectivity meta-analyses. Brain Imaging Behav. 16, 1538–1551 (2022).

63. Esser, S. K., Hill, S. & Tononi, G. Breakdown of Effective Connectivity During Slow Wave Sleep: Investigating the Mechanism Underlying a Cortical Gate Using Large-Scale Modeling. J. Neurophysiol. 102, 2096–2111 (2009).

64. Massimini, M. et al. Breakdown of Cortical Effective Connectivity During Sleep. Science 309, 2228–2232 (2005).

65. Marek, S. et al. Reproducible brain-wide association studies require thousands of individuals. Nature 603, 654–660 (2022).

66. Waskom, M. L. Seaborn: statistical data visualization. J. Open Source Softw. 6, 3021 (2021).

67. Fischl, B. FreeSurfer. NeuroImage 62, 774–781 (2012).

68. Hariri, A. R., Tessitore, A., Mattay, V. S., Fera, F. & Weinberger, D. R. The Amygdala Response to Emotional Stimuli: A Comparison of Faces and Scenes. NeuroImage 17, 317– 323 (2002).

69. Barch, D. M. et al. Function in the human connectome: Task-fMRI and individual differences in behavior. NeuroImage 80, 169–189 (2013).

70. Kriegeskorte, N., Mur, M. & Bandettini, P. Representational similarity analysis – connecting the branches of systems neuroscience. Front. Syst. Neurosci. 2, (2008).

71. Jenkinson, M., Beckmann, C. F., Behrens, T. E. J., Woolrich, M. W. & Smith, S. M. FSL. NeuroImage 62, 782–790 (2012).

72. Glasser, M. F. et al. The minimal preprocessing pipelines for the Human Connectome Project. NeuroImage 80, 105–124 (2013).

