## Supplementary info for "Sleep-insomnia superposition: opposing brain signatures of sleep in task-based and resting-state conditions"

Daniel Felsky PhD

Independent Scientist, Krembil Centre for Neuroinformatics, Centre for Addiction and Mental Health

Assistant Professor, Department of Psychiatry and Dalla Lana School of Public Health, University of Toronto

12th Floor, 250 College Street, Toronto ON, M5T 1R8, Canada

www.felskylab.com

(416) 535 8501 x33587

### Supplementary Information

#### Supplementary Results

##### Task fMRI decoding results

We were able to successfully decode this binary condition using signal from all brain regions including non-visual brain regions; the success of classifiers from non-visual regions may be attributable to the ease of the decoding task. It could also be attributed to expectation signals where the participant is anticipating the next stimulus, a process that involves many non-visual regions across the brain[1]. Association models revealed that the superior parietal regions were those linked to all three of duration of longest sleep bout, depressive symptoms, and cognition. This region is a multimodal integration area that receives input from the dorsal stream[2] and is responsible for cue integration, spatial orientation, and motor movement planning[3–9]. The triangulation of this region as highly important is reasonable since the visual task involved motor action (button pressing) making a selection based on spatial location of the matching stimulus. This means that superior parietal regions would be downstream of the processing pathway for performing this task.

Other phenotypes showed associations with multivariate neural coding. Cognition showed associations that spanned across the whole visual cortex except for some intermediate regions including early visual areas and areas along both the ventral and dorsal pathways. It also showed a positive association with large regions in the frontal lobe as well. This indicates that cognitive ability is reflected along both sensory and cognitive regions in line with most of the existing literature about cognitive ability[10,11]. Depressive symptoms had associations across the whole brain with higher depressive symptoms being associated with lower multivariate neural activity. The size of this effect is larger than what was reported by most of the previous studies[12,13] which could be attributed to the large sample size utilized here leading to more statistical power. Self-reported insomnia and daytime dozing had almost no significant associations but the brain-wide association patterns showed high correlation with depressive symptoms.

#### Resting-state connectivity between default mode network and motor cortex is associated with insomnia and depression

Functional connectivity data also showed an association of both self-reported insomnia and depressive symptoms and the connectivity default mode network component (IC12) and the somatomotor network component (IC10). Zooming into these results using the seed-based connectivity analysis, we found increased connectivity from the regions STSva and SFL from IC12 to the areas 1, 2, 3a, and 3b from the motor cortex and areas Ig, OP4, and PoI2 from the opercular cortex (**Figure S6**). Area STSva is a part of the auditory association cortex though it is believed to be more multimodal [14] while area SFL is known as a predominantly language area[14,15]. Areas 1,2, 3a, and 3b are all part of the primary somatosensory network. Region 4 which is the primary motor cortex did not show a significant association of connectivity meaning that the effect was restricted to somatosensory areas. Areas Ig, OP4, and PoI2 are part of the opercular and insular cortices. These regions are also somatosensory that were associated with sensation in the mouth and pharynx areas[16] and pain sensation[17]. Pain sensation has been associated with depressive symptoms in multiple accounts[18–20] including a lowering of pain threshold as a result of depression[21,22]. Increased frequency of self-reported insomnia also additonally associated with a significantly increased interconnectivity between regions of the IC12 while increased depressive symptoms was associated with a decreased connectivity withing the somatosensory areas in IC10.

#### Early visual cortex exhibits cortical thickening with depression, lack of sleep, and lower cognitive performance

We built OLS models to measure associations of cortical thickness with the phenotypes of interest in regions defined according to the HCP parcellation. Measured duration of longest sleep bout, depression score, and cognition all showed diffuse significant associations but with a considerable overlap along the auditory, insular, and temporal regions (**Figure S7A**). Measured duration of longest sleep bout was found to have a widespread association with additional associations across the paracentral, frontal, and motor regions (**Figure S7B**). The association was positive with longer continuous sleep being associated with higher cortical thickness in almost all brain regions except for the primary visual cortex (V1) and early visual cortex (V2). Self-reported insomnia and daytime dozing did not show any significant difference in cortical thickness values. PHQ-2 score also showed sparsely-distributed significant associations with higher depression scores associated with cortical thinning especially in the temporal, motor, and medial frontal regions. The invested association of V1 persisted also in the depression score. Cognition showed significant associations in the temporal and frontal regions with higher score being associated with higher cortical thickness except for the areas in the early visual cortex V1, V2, and V4. Correlations between the associations of anatomical models were largely consistent with those from the task fMRI experiment. Measure duration of longest sleep bout correlated positively with cognitive score and negatively with depression score, frequency of insomnia, and frequency of daytime dozing. Depression score coefficients correlated positively with frequency of insomnia and daytime dozing but negatively with cognition. Cognition, correlated negatively with both  frequency of insomnia and daytime dozing though the former was a weak anti-correlation. Self-reported frequencies of insomnia and daytime dozing correlated positively.

These results are largely similar patterns to the task fMRI data where duration of longest sleep bout associations were similar to those of daytime dozing and opposite to those of insomnia and depressive symptoms but with  significant overlap at the insular, frontal, and temporal regions and at the primary and early visual areas. It shows a counter-intuitive result where lower quality sleep is associated with thicker V1 and V2. This could be explained as a compensation mechanism for impaired visual processing due to low-quality sleep[23]. The causal relationship could also be reversed where a thicker V1/2 is causing a more sensitive visual system which in turn causes lower sleep quality.These results were observed in other conditions in mostly developmental studies where it was suggested that thinner cortex could be associated with more synaptic pruning leading to a better optimized visual cortex[24,25].

Supplementary tables

**Table S1**: Statistical summary of the participants included in the study and the co-variates used in the linear models.

| Variable | Female | Male |
|---|---|---|
| Count | 16497 (53.3%) | 14448 (46.7%) |
| Age mean (SD) | 62.7 (7.3) | 64.0 (7.6) |
| Socioeconomic status (Townsend deprivation index) | -1.90 (2.69) | -1.97 (2.70) |
| Education level (Age completed full time | 16.87 (1.50) | 16.88 (1.69) |

| | | |
|---|---|---|
| education) | | |
| Ethnic background | White 16037 (97.4%)<br>Asian or Asian British 103 (0.63%)<br>Black or Black British 96 (0.58%)<br>Mixed 90 (0.55%)<br>Other 85 (0.52%)<br>Chinese 47 (0.29%) | White 13972 (97.0%)<br>Asian or Asian British 191 (1.33%)<br>Black or Black British 84 (0.58%)<br>Mixed 68 (0.47%)<br>Other 48 (0.33%)<br>Chinese 34 (0.24%) |
| Difference between MRI acquisition and Actigraphy mean (SD) | 2.6 (1.7) | 2.6 (1.7) |
| Scan site | Cheadle 10373 (62.9%)<br>Newcastle 4170 (25.3%)<br>Reading 1954 (11.8%) | Cheadle 9385 (65.0%)<br>Newcastle 3403 (23.6%)<br>Reading 1660 (11.5%) |
| Head motion (resting) mm | 0.11 (0.05) | 0.13 (0.06) |
| Head motion (task) mm | 0.14 (0.06) | 0.15 (0.06) |
| Mean task accuracy % | 0.89 (0.19) | 0.90 (0.18) |
| Task response time mean ms | 877.8 (201.2) | 884.7 (199.8) |
| Task response time standard deviation ms | 323.6 (0.6.1) | 315.0 (104.8) |

**Table S2**: Statistical summary of the phenotypes measured and percentage of missing values from the overall cohort from table S1

| Variable | | Female | Male |
|---|---|---|---|
| Duration of longest sleep bout (measured by accelerometer) | Count | 7145 (55.0%) | 5848 (45%) |
| | Missingness % | 56.7% | 59.5% |
| | Mean (SD) | 6.78 (1.49) | 6.22 (1.65) |
| PHQ-2 score | Count | 15751 (53.1%) | 13899 (46.9%) |
| | Missingness % | 4.52% | 3.80% |
| | Mean (SD) | 4.44 (0.96) | 4.36 (0.85) |
| Sleeplessness/Insomnia (self-reported) | Count | 16347 (53.3%) | 14310 (46.7%) |
| | Missingness % | 0.91% | 0.96% |
| | Mean (SD) | 2.19 (0.69) | 1.99 (0.75) |
| Cognition score (Symbol-digit substitution task) | Count | 10353 (53.3%) | 9062 (46.7%) |
| | Missingness % | 37.2% | 37.3% |
| | Mean (SD) | 19.33 (5.28) | 18.88 (5.15) |
| Daytime dozing (self- | Count | 16339 | 14303 |

| | | | |
|---|---|---|---|
| reported) | Missingness | 0.96% | 1.00% |
| | Mean (SD) | 2.22 (0.46) | 2.28 (0.50) |

**Table S3**: UK-Biobank variable codes

| Variable | Code | Instance |
|---|---|---|
| Age | 21022 | 0 |
| Date of attending assessment centre | 53 | 0, 2 |
| Sex at birth | 31 | 0 |
| Socioeconomic status (Townsend deprivation index) | 189 | 0 |
| Education level | 845 | 0 |
| Ethnic background | 21000 | 0 |
| Imaging site | 54 | 2 |
| Task fMRI brain images | 20249 | 2 |
| Resting fMRI brain images | 20227 | 2 |
| Functional connectivity (full | 25750 | 2 |

| | | |
|---|---|---|
| correlation function, dimension 25) | | |
| Brain anatomy (Freesurfer) | 20263 | 2 |
| Mean task fMRI head motion | 25742 | 2 |
| Mean resting fMRI head motion | 25741 | 2 |
| Task fMRI behavioral parameters | 25749 | 2 |
| Cognition (Number of symbol digit matches made correctly, Symbol-digit substitution task) | 23324 | 2 |
| PHQ-2 (Frequency of depressed mood in last 2 weeks) | 2050 | 2 |
| PHQ-2 (Frequency of unenthusiasm / disinterest in last 2 weeks) | 2060 | 2 |
| Sleeplessness / insomnia | 1200 | 2 |
| Daytime dozing / sleeping (narcolepsy) | 1220 | 2 |

**Table S4:** Resting-state correlation value comparison between Fan et al. and our results (* indicates that Fan et al. results are from correlation with self-reported sleep duration while ours are from accelerometer duration of longest sleep bout)

| Phenotype pair | Fan el al. results correlation | Our results |
|---|---|---|
| Insomnia-daytime dozing | -0.66 | -0.59 |
| Insomnia-sleep duration* | 0.59 | 0.51 |
| Daytime dozing-sleep duration* | -0.86 | -0.84 |

Supplementary figures

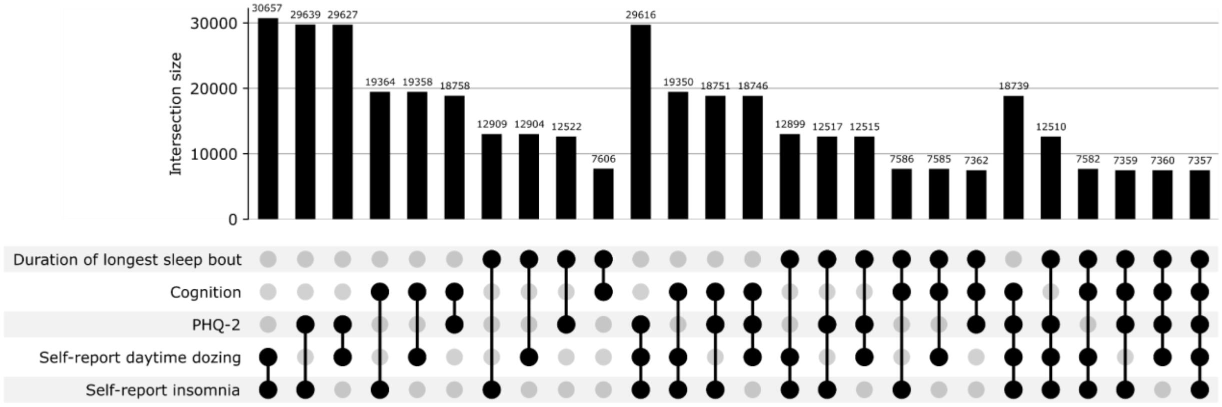


**Figure S1**: Upset plot showing the overlap between existing measurements of the different phenotypes of interest.

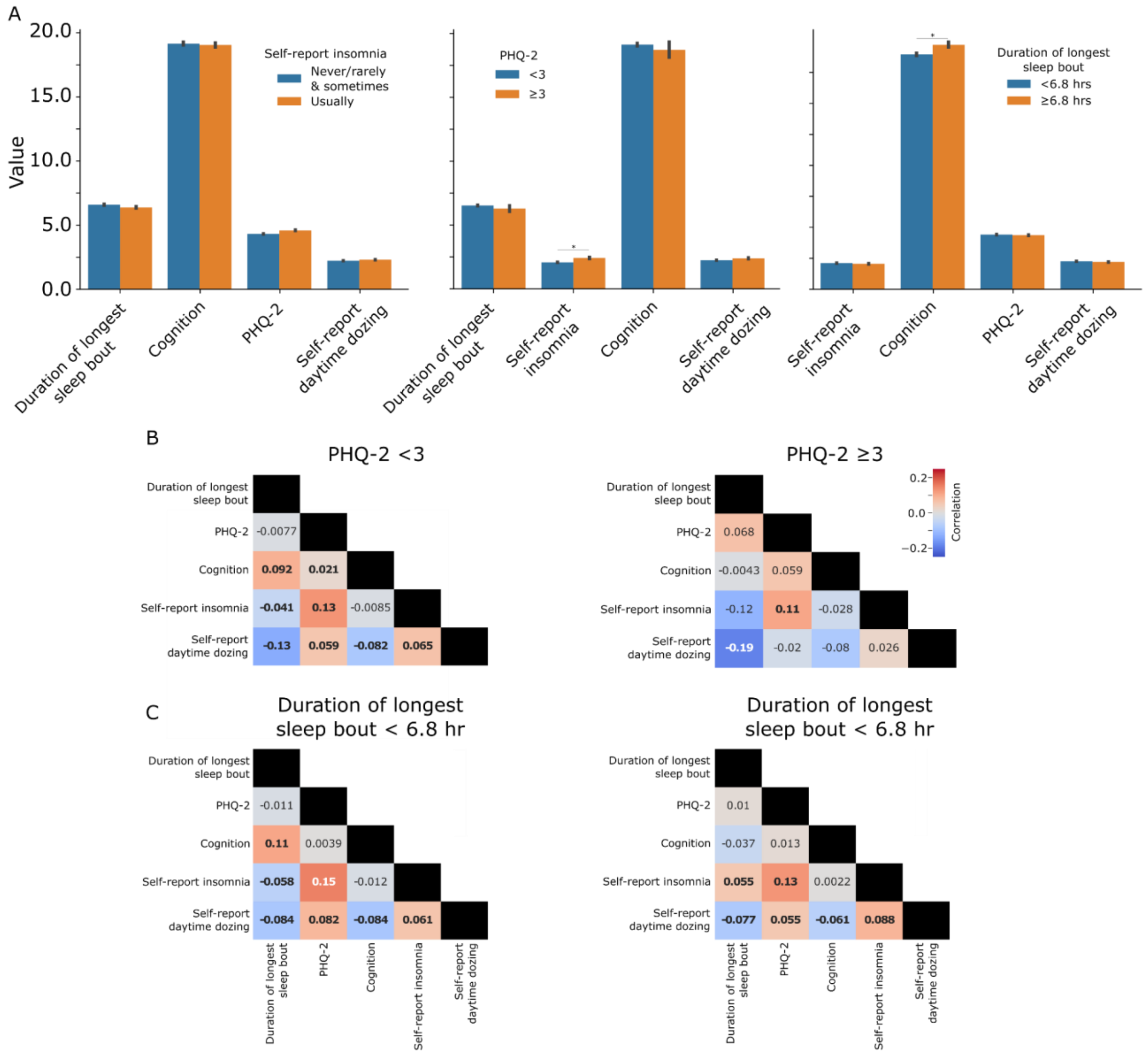


**Figure S2**: Statistical description of the phenotypes grouped by different factors. **A** shows mean values of different phenotypes split by the self-reporting of insomnia, PHQ-2, and duration of longest sleep bout. **B** Pairwise correlation between the phenotypes when split by PHQ-2 score representing the difference between depressed and non-depressed groups. **C** Pairwise correlation between the phenotypes when split by duration of longest sleep bout by the median value of 6.8 hours. Bolded values are statistically significant.

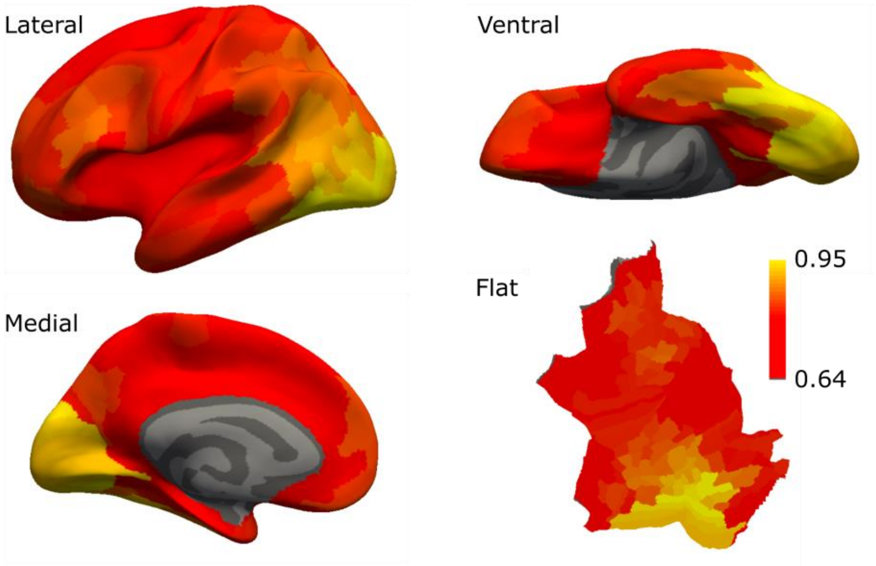


**Figure S3**: fMRI decoding was successful from all brain regions with higher accuracy in visual regions. Brain maps show mean decoding accuracy from all brain regions in different views.

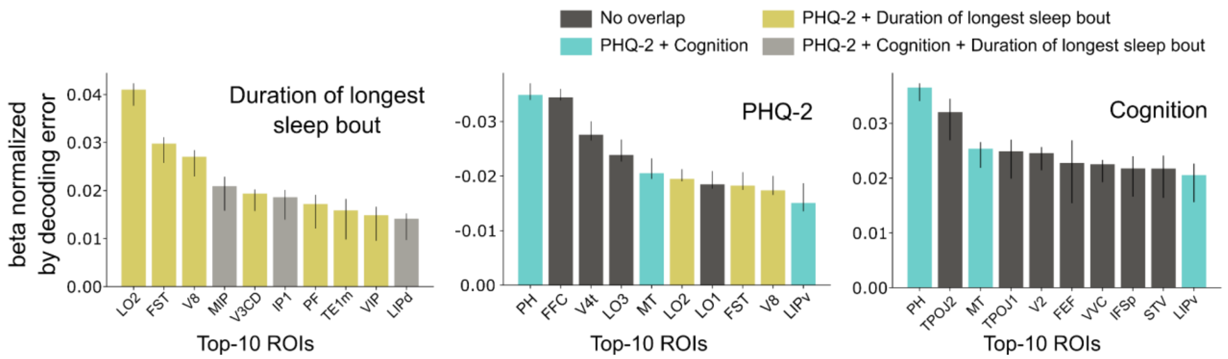


**Figure S4**: Top 10 significant coefficients normalized by decoding error for each of the phenotypes that had 10 or more significant regions color-coded to show the overlapping phenotypes following the code in C.

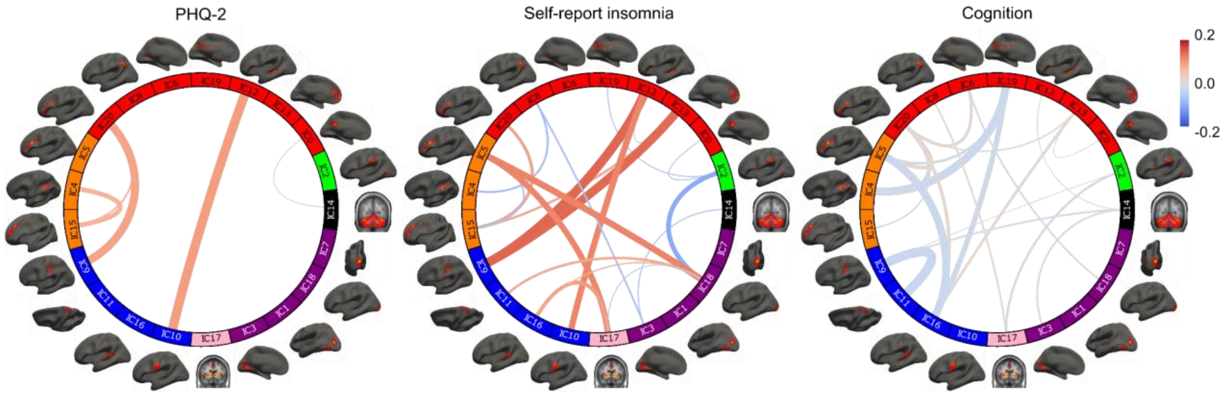


**Figure S5**: Resting-state connectivity association results with the remaining phenotypes (PHQ-2 score, self-reported insomnia, and cognition test score).

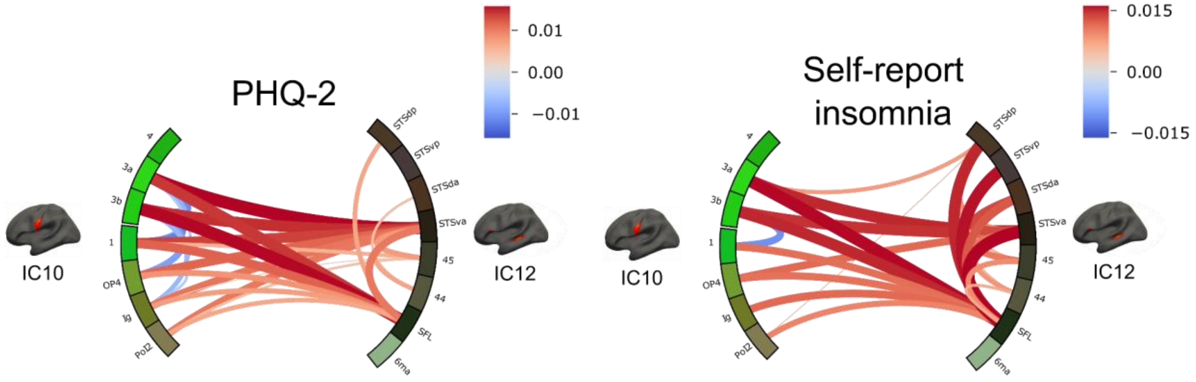


**Figure S6**: Seed-based correlation associations results between IC12 and IC10 regions.

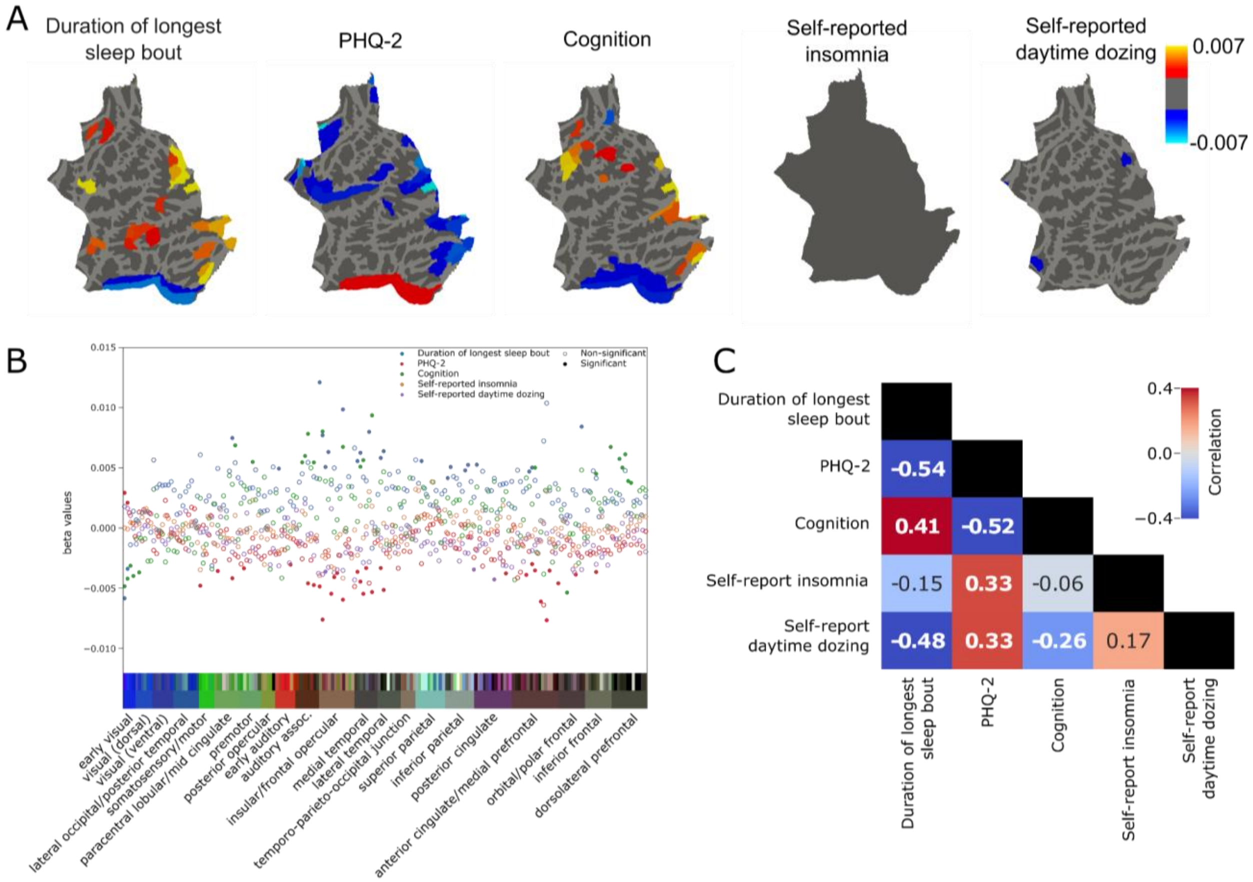


**Figure S7**: Results of associations of cortical thickness with the different phenotypes. **A** shows the region maps with significant associations color coded by the beta values for each of the phenotype models. **B** summarizes the overall beta values of the models. The regions are organized and color coded according to their groupings in the human connectome project [14]**. C** shows the pairwise correlation values between coefficients from each phenotype model across all regions.

### References


1. Gilbert, C. D. & Li, W. Top-down influences on visual processing. *Nat. Rev. Neurosci.* **14**, 10.1038/nrn3476 (2013).
2. Alahmadi, A. A. S. Investigating the sub-regions of the superior parietal cortex using functional magnetic resonance imaging connectivity. *Insights Imaging* **12**, 47 (2021).
3. Bray, S., Arnold, A. E. G. F., Iaria, G. & MacQueen, G. Structural connectivity of visuotopic intraparietal sulcus. *NeuroImage* **82**, 137–145 (2013).
4. Corbetta, M., Shulman, G. L., Miezin, F. M. & Petersen, S. E. Superior Parietal Cortex Activation During Spatial Attention Shifts and Visual Feature Conjunction. *Science* **270**, 802–805 (1995).
5. Culham, J. C. & Valyear, K. F. Human parietal cortex in action. *Curr. Opin. Neurobiol.* **16**, 205–212 (2006).
6. Lloyd, D., Morrison, I. & Roberts, N. Role for Human Posterior Parietal Cortex in Visual Processing of Aversive Objects in Peripersonal Space. *J. Neurophysiol.* **95**, 205–214 (2006).
7. Wang, J. *et al.* Convergent functional architecture of the superior parietal lobule unraveled with multimodal neuroimaging approaches. *Hum. Brain Mapp.* **36**, 238–257 (2015).
8. Weiss, P. H., Marshall, J. C., Zilles, K. & Fink, G. R. Are action and perception in near and far space additive or interactive factors? *NeuroImage* **18**, 837–846 (2003).
9. Wenderoth, N., Debaere, F., Sunaert, S., Hecke, P. van & Swinnen, S. P. Parieto-premotor Areas Mediate Directional Interference During Bimanual Movements. *Cereb. Cortex* **14**, 1153–1163 (2004).
10. Eyler, L. T., Sherzai, A., Kaup, A. R. & Jeste, D. V. A Review of Functional Brain Imaging Correlates of Successful Cognitive Aging. *Biol. Psychiatry* **70**, 115–122 (2011).
11. Altena, E. *et al.* Prefrontal hypoactivation and recovery in insomnia. *Sleep* **31**, 1271–1276

(2008).

12. Palmer, S. M., Crewther, S. G., Carey, L. M. & The START Project Team. A Meta-Analysis of Changes in Brain Activity in Clinical Depression. *Front. Hum. Neurosci.* **8**, (2015).
13. Desseilles, M. *et al.* Abnormal Neural Filtering of Irrelevant Visual Information in Depression. *J. Neurosci.* **29**, 1395–1403 (2009).
14. Glasser, M. F. *et al.* A multi-modal parcellation of human cerebral cortex. *Nature* **536**, 171–178 (2016).
15. Fujii, M. *et al.* Intraoperative subcortical mapping of a language-associated deep frontal tract connecting the superior frontal gyrus to Broca's area in the dominant hemisphere of patients with glioma. *J. Neurosurg.* **122**, 1390–1396 (2015).
16. Mălîia, M.-D. *et al.* Functional mapping and effective connectivity of the human operculum. *Cortex J. Devoted Study Nerv. Syst. Behav.* **109**, 303–321 (2018).
17. Garcia-Larrea, L. The posterior insular-opercular region and the search of a primary cortex for pain. *Neurophysiol. Clin. Neurophysiol.* **42**, 299–313 (2012).
18. Stahl, S. & Briley, M. Understanding pain in depression. *Hum. Psychopharmacol. Clin. Exp.* **19**, S9–S13 (2004).
19. Fields, H. Depression and Pain A Neurobiological Model. *Cogn. Behav. Neurol.* **4**, 83–92 (1991).
20. Korff, M. V. & Simon, G. The Relationship Between Pain and Depression. *Br. J. Psychiatry* **168**, 101–108 (1996).
21. Marazziti, D. *et al.* Pain threshold is reduced in depression. *Int. J. Neuropsychopharmacol.* **1**, 45–48 (1998).
22. Klauenberg, S. *et al.* Depression and changed pain perception: Hints for a central disinhibition mechanism. *PAIN* **140**, 332–343 (2008).
23. Burge, W. K. *et al.* Cortical thickness in human V1 associated with central vision loss. *Sci. Rep.* **6**, 23268 (2016).

24. Bhat, A., Biagi, L., Cioni, G., Tinelli, F. & Morrone, M. C. Cortical thickness of primary visual cortex correlates with motion deficits in periventricular leukomalacia. *Neuropsychologia* **151**, 107717 (2021).
25. Jiang, J. *et al.* Thick Visual Cortex in the Early Blind. *J. Neurosci.* **29**, 2205–2211 (2009).
